# The *Drosophila* DCP2 is evolutionarily conserved in sequence and structure – insights from *in silico* studies of DmDCP2 orthologs and paralogs

**DOI:** 10.1101/2021.04.18.440350

**Authors:** Rohit Kunar, Jagat Kumar Roy

## Abstract

The mRNA decapping proteins (DCPs) function to hydrolyze the 7-methylguanosine cap at the 5’ end of mRNAs thereby, exposing the transcript for degradation by the exonuclease(s) and hence, play a pioneering role in the mRNA decay pathway. In *Drosophila melanogaster*, the mRNA decapping protein 2 (DCP2) is the only catalytically active mRNA decapping enzyme present. Despite its presence being reported across diverse species in the phylogenetic tree, a quantitative approach to the index of its conservation in terms of its sequence has not been reported so far. With structural and mechanistic insights being explored in the yeasts, the insect DCP2 has never been explored in the perspectives of structure and the indices of the conservation of its sequence and/or structure *vis-à-vis* topological facets. Being an evolutionarily conserved protein, the present endeavor aimed at deciphering the evolutionary relationship(s) and the pattern of conservation of the sequence of DCP2 across the phylogenetic tree as well as in sibling species of *D. melanogaster* through a semi-quantitative approach relying on multiple sequence alignment and analyses of percentage identity matrices. Since NUDIX proteins are functionally diverse, an attempt to identify the other NUDIX proteins (or, DCP2 paralogs) in *D. melanogaster* and compare and align their structural features with that of DCP2 through *in silico* approaches was endeavored in parallel. Our observations provide quantitative and structural bases for the observed evolutionary conservation of DCP2 across the diverse phyla and also, identify and reinforce the structural conservation of the NUDIX family in *D. melanogaster*.

## Introduction

mRNA decapping, by virtue of its pioneering role in the degradation of transcripts plays a significant role in the turnover of mRNA and widely affects the expression of genes (Mitchell and Tollervey, 2001; Raghavan and Bohjanen, 2004; Song et al, 2010). The mRNA decapping protein 2 (DCP2) performs this essential step (Dunckley and Parker, 1999) and is conserved across diverse species (Wang et al, 2002). Previous studies (reviewed in Grudzien-Nogalska and Kiledjian, 2016; Wurm and Sprangers, 2019) have identified DCP2 to contain a Box A domain and a NUDIX motif. The Box A domain is essential to maintain the catalytic fidelity of decapping (She et al, 2008) and interacts with the activator of decapping, the mRNA decapping protein 1 (DCP1) (Li and Kiledjian, 2010). Another domain, the Box B domain, which is a part of the NUDIX fold and lies just C-terminal to the NUDIX motif, is essential for binding to the RNA (She et al, 2008).

The NUDIX hydrolase superfamily or as commonly referred to as the homology clan (Srouji et al, 2017) encompasses approximately 80,000 proteins, which bears the 23 residue consensus sequence, GX_5_EX_7_REUXEEXGU (U: bulky hydrophobic aliphatic residue such as leucine, isoleucine or valine; X: any amino acid) and harbor a helix-loop-helix structure. The proteins are characterized by the presence of a beta-grasp domain composed of approximately 130 residues which coordinates Mg^2+^ ions *via* three conserved Glutamate residues. Members of this superfamily, show monophyly with regard to their function and belong to four general functional classes, *viz*., pyrophosphohydrolases, isopentenyl diphosphate isomerases (IDIs), adenine/guanine (A/G) mismatch-specific adenine glycosylases, and some proteins with non-enzymatic activities such as protein-protein interaction and/or transcriptional regulation. The largest sub-group, pyrophosphohydrolases encompass proteins with more than hundred distinct hydrolase specificities (Srouji et al., 2017).

During evolution, protein structures change due to mutations, which can involve substitutions or insertions or deletions. The extent of perturbations in the structures of higher order in a protein depend on the type of mutations, where, some mutations may completely disrupt the existing structure while others may affect the physic-chemical properties of the protein (and confer altered or neo-functional properties) without causing major perturbations of the structure (Illergård et al, 2009). Protein domains are the modules of protein architecture which are composed of independently folding subsequences which are found to be arranged differently in different proteins (Forslund et al, 2011). Usually, the conservation of homologous sequences is rarely homogeneous along their length, with their conservation being localized to specific regions. Characteristically, the most conserved regions in a protein are those which are important functionally and structurally (Sitbon and Pietrokovski, 2007).

In perspective, the *Drosophila* DCP2 is orthologous to the isoform(s) harbored by all other species in the phylogenetic tree and is paralogous to the different proteins harboring the NUDIX motif *vis-à-vis* NUDIX proteins in *Drosophila* itself (Jensen, 2001). Herein, the index of homology of the sequence of the various DCP2 orthologs and paralogs has been explored alongwith insights into the structure of *Drosophila* DCP2 and its homology with the paralogous proteins.

## Materials and Methods

### Retrieval of sequences of DCP2 orthologs and paralogs

Sequences of DCP2 orthologs across the phylogenetic tree and sibling species of *Drosophila melanogaster* were procured from the NCBI (http://www.ncbi.nlm.nih.gov), *viz. Disctyostelium discoideum* (XP_639160.2), *Saccharomyces cerevisiae* (KZV08504.1), *Schizosaccharomyces pombe* (NP_593780.1), *Caenorhabditis elegans* (NP_502609.2), *Strongylocentrotus purpuratus* (XM_781343.4), *Danio rerio* (AAH66577.1), *Xenopus tropicalis* (CAJ83772.1), *Gallus gallus* (XP_004949329.1), *Mus musculus* (NP_081766.1), *Rattus norwegicus* (NP_001163940.1), *Homo sapiens* (EAW48988.1), *Drosophila melanogaster* (NP_648805.2), *D. ananassae* (XP_001957344.1), *D. erecta* (XP_015013047.1), *D. grimshawi* (XP_001985489.1), *D. mojavensis* (XP_002008663.2), *D. persimilis* (XP_002021219.1), *D. pseudoobscura pseudoobscura* (XP_001353423.3), *D. sechellia* (XP_002030644.1), *D. simulans* (XP_016032029.1), *D. virilis* (XP_002048412.1), *D. willistoni* (XP_023031850.1) and *D. yakuba* (XP_015045072.1).

Sequences of the DCP2 paralogs in *Drosophila melanogaster* were procured from the FlyBase (http://www.flybase.org), *viz*., CG2091 (NP_649582), Apf (NP_723505), Aps (NP_648421), CG8128 (NP_573053), CG12567 (NP_001015384), CG42813 (NP_733202), CG10898 (CG_650083), CG18094 (NP_609974), CG10194 (NP_609973), CG10195 (NP_609970), CG11095 (NP_572927) and CG42814 (NP_001189301).

The sequences were manually analysed for sequence features, *viz*., amino acid sequence and size of the Box A, spacer and NUDIX domains and the remaining N- and C-terminal regions, and a graphical representation of the same was performed using MS-Excel 2010.

### Multiple Sequence Alignment (MSA), Percent Identity Matrix (PIM) analyses and evolutionary relationships

Multiple Sequence Alignment (MSA) and calculation of Percent Identity Matrix (PIM) were performed using Clustal Omega (EBI; https://www.ebi.ac.uk/Tools/msa/clustalo/), while evolutionary relationships were inferred using the Maximum Likelihood method using the PhyML (Guindon et al., 2010) software (http://www.atgc-montpellier.fr/phyml/).

**Structural analyses**

The secondary structure of the DCP2-PA isoform was deduced using PSIPRED (Jones, 1999; http://bioinf.cs.ucl.ac.uk/psipred/) tool, while the tertiary structure was determined using the I-TASSER (Roy *et al*., 2010; https://zhanglab.ccmb.med.umich.edu/I-TASSER/) tool. The structure was visualized and analysed for the different features with an offline tool, UCSF Chimera (NIH; https://www.cgl.ucsf.edu/chimera/). The backbone confirmation was evaluated by inspection of the Phi/Psi angles using the Ramachandran plot (Ramachandran *et al*., 1963) using the RAMPAGE tool, available online (http://mordred.bioc.cam.ac.uk/∼rapper/rampage.php).

## Results

### The sequence of NUDIX domain of DCP2 is much more conserved than the rest of the protein sequence

The linear sequence of amino acids in DCP2 can be conveniently categorized into five different segments, *viz*., the N-terminal region, *i*.*e*., from the pioneering amino acid to the Box A domain, the Box A domain, the spacer tripeptide, the NUDIX domain and the C-terminal region, *i*.*e*., after the NUDIX domain till the carboxy-terminus. The orthologs were first analysed for the total number of amino acids and further assessed for the number of amino acids harbored in or constituting the individual segments. Since the Box A and the NUDIX domains are the two classic domains of DCP2, they along with the spacer tripeptide were analysed for the identity of the pioneering and terminal amino acid residues. The proteins were then assessed for the conservation of sequence of the complete protein and then for the NUDIX domain only, following which, the molecular phylogenetic relationships were also identified.

#### a. Vertebrate orthologs of DCP2 show higher degree of conservation than invertebrate counterparts

Among the orthologs harbored by the representative species of the phylogenetic tree, except for *Dictyostelium, Caenorhabditis* and *Drosophila*, all other species have a short N-terminal or pre-Box A sequence and harbor the two classic domains close to the N-terminal. The Box A domain does not vary appreciably in length (82-85 residues), except for the *Drosophila melanogaster* ortholog, wherein it is 95 residues long. While the pioneering residue of the Box A domain varies in the invertebrates, the C-terminal residue is conserved in all the species {Tyrosine (Y)}. The Box A domain is followed by the spacer sequence, which is a tripeptide stretch and begins with Lysine (K). The lower members show variability in composition but the vertebrates show a constant conserved sequence of Lysine (K) – Methionine (M) – Serine (S). The NUDIX domain is conserved in length and sequence and almost always starts with Valine (V), except for the yeasts, *Saccharomyces cerevisiae* and *Schizosaccharomyces pombe*, wherein it starts with Isoleucine (I). The terminal residue is variable however, but is conserved in the vertebrates. Notably, the vertebrate orthologs are shorter in length as compared to the lower members, and show a higher degree of conservation, both in composition and location of sequence and in size. Moreover, the number of amino acids constituting the protein decreases in the vertebrate phyla (**Table 1 and Figure 1**). Closer analyses of the complete DCP2 sequence and the sequence of the NUDIX motif harbored, through percentage identity matrices (**Figure 2 A and B**), shows that the vertebrate orthologs are more conserved in either case as compared to their invertebrate predecessors. Strikingly, although the two yeast species analysed, occupy similar positions in the evolutionary tree, they show extremely low homology among themselves. Moreover, the *S. cerevisiae* (budding yeast) ortholog shows the least homology with all the other species analysed. However, in all the species considered here, the sequence of the NUDIX domain shows a better index of homology as compared to the sequence of the entire DCP2 protein.

**Table 1:**
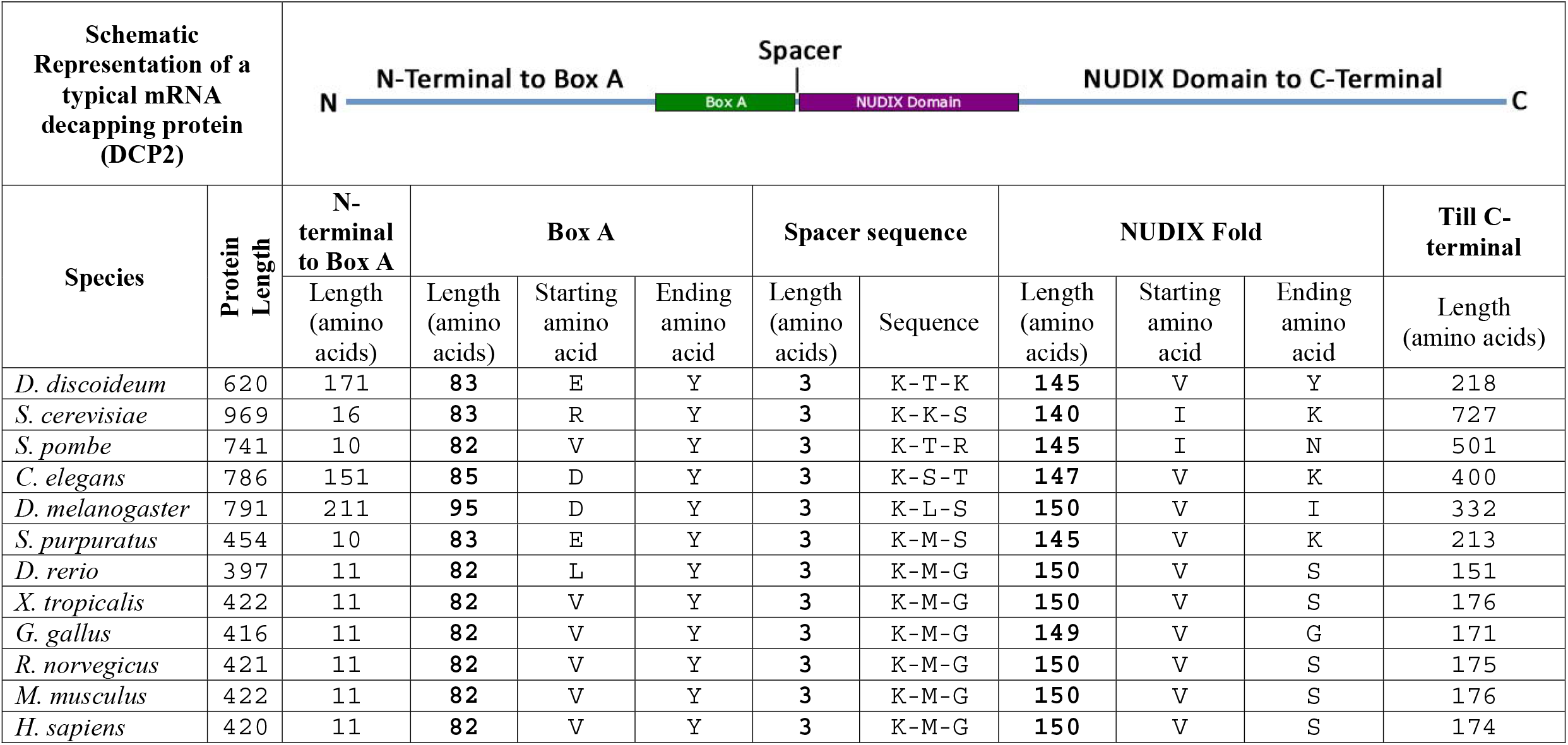
Table showing the sizes of the different regions of the DCP2 orthologs across the phylogenetic tree. The NUDIX domain is conserved in length and sequence and almost always starts with Valine (V), except for *Saccharomyces cerevisiae* and *Schizosaccharomyces pombe*, which start with Isoleucine (I). The terminal residue is variable however, but is conserved in the vertebrates. The Box A domain does not vary appreciably in length, except for *Drosophila melanogaster* which is 95 residues long. The C-terminal residue however, is Tyrosine (Y) across the species, and is conserved. The Box A domain is followed by the spacer sequence towards the C-terminal, which is three amino acids long and begins with Lysine (K). The lower members show variability in composition but the vertebrates show a constant conserved sequence of Lysine(K)–Methionine(M)–Serine(S). Notably, the vertebrate orthologs are shorter in length as compared to the lower members, and show a higher degree of conservation, both in composition and location of sequence and in size.

| Schematic Representation of a typical mRNA decapping protein (DCP2) |  |  |  |  |  |  |  |  |  |  |  |
| --- | --- | --- | --- | --- | --- | --- | --- | --- | --- | --- | --- |
| Species | Protein Length | N-terminal to Box A | Box A |  |  | Spacer sequence |  | NUDIX Fold |  |  | Till C-terminal |
|  |  | Length (amino acids) | Length (amino acids) | Starting amino acid | Ending amino acid | Length (amino acids) | Sequence | Length (amino acids) | Starting amino acid | Ending amino acid | Length (amino acids) |
| *D. discoideum* | 620 | 171 | 83 | E | Y | 3 | K-T-K | 145 | V | Y | 218 |
| *S. cerevisiae* | 969 | 16 | 83 | R | Y | 3 | K-K-S | 140 | I | K | 727 |
| *S. pombe* | 741 | 10 | 82 | V | Y | 3 | K-T-R | 145 | I | N | 501 |
| *C. elegans* | 786 | 151 | 85 | D | Y | 3 | K-S-T | 147 | V | K | 400 |
| *D. melanogaster* | 791 | 211 | 95 | D | Y | 3 | K-L-S | 150 | V | I | 332 |
| *S. purpuratus* | 454 | 10 | 83 | E | Y | 3 | K-M-S | 145 | V | K | 213 |
| *D. rerio* | 397 | 11 | 82 | L | Y | 3 | K-M-G | 150 | V | S | 151 |
| *X. tropicalis* | 422 | 11 | 82 | V | Y | 3 | K-M-G | 150 | V | S | 176 |
| *G. gallus* | 416 | 11 | 82 | V | Y | 3 | K-M-G | 149 | V | G | 171 |
| *R. norvegicus* | 421 | 11 | 82 | V | Y | 3 | K-M-G | 150 | V | S | 175 |
| *M. musculus* | 422 | 11 | 82 | V | Y | 3 | K-M-G | 150 | V | S | 176 |
| *H. sapiens* | 420 | 11 | 82 | V | Y | 3 | K-M-G | 150 | V | S | 174 |

**Figure 1:**
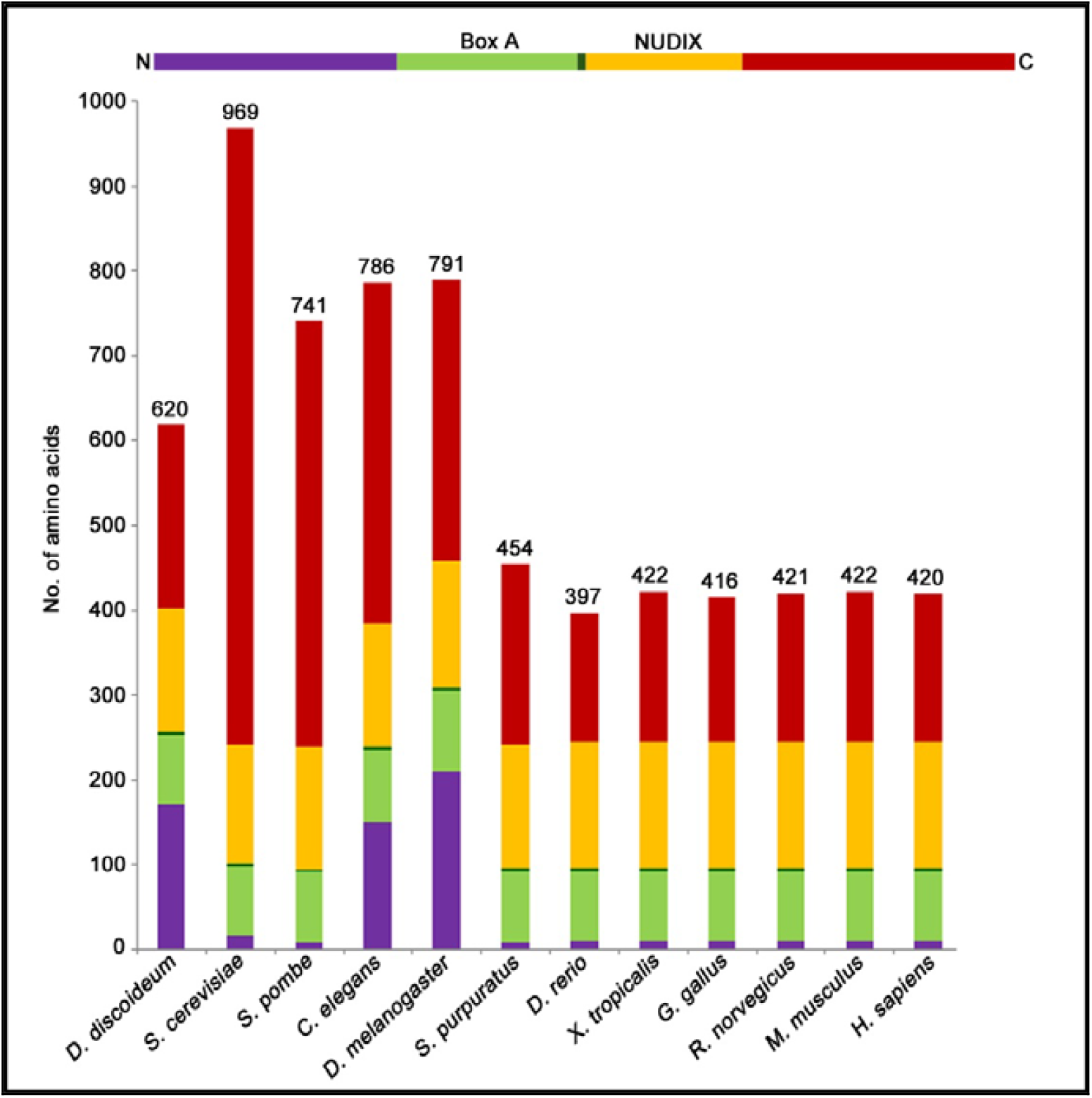
Graphical representation of the different regions of the DCP2 orthologs across the phylogenetic tree. The representative invertebrate orthologs harbor longer sequences while the representative vertebrate orthologs comprise of lesser number of residues. While *Saccharomyces cerevisiae* harbors the longest ortholog, the vertebrate orthologs do not show appreciable difference in the total number of amino acids as well as those constituting the individual regions.

**Figure 2:**
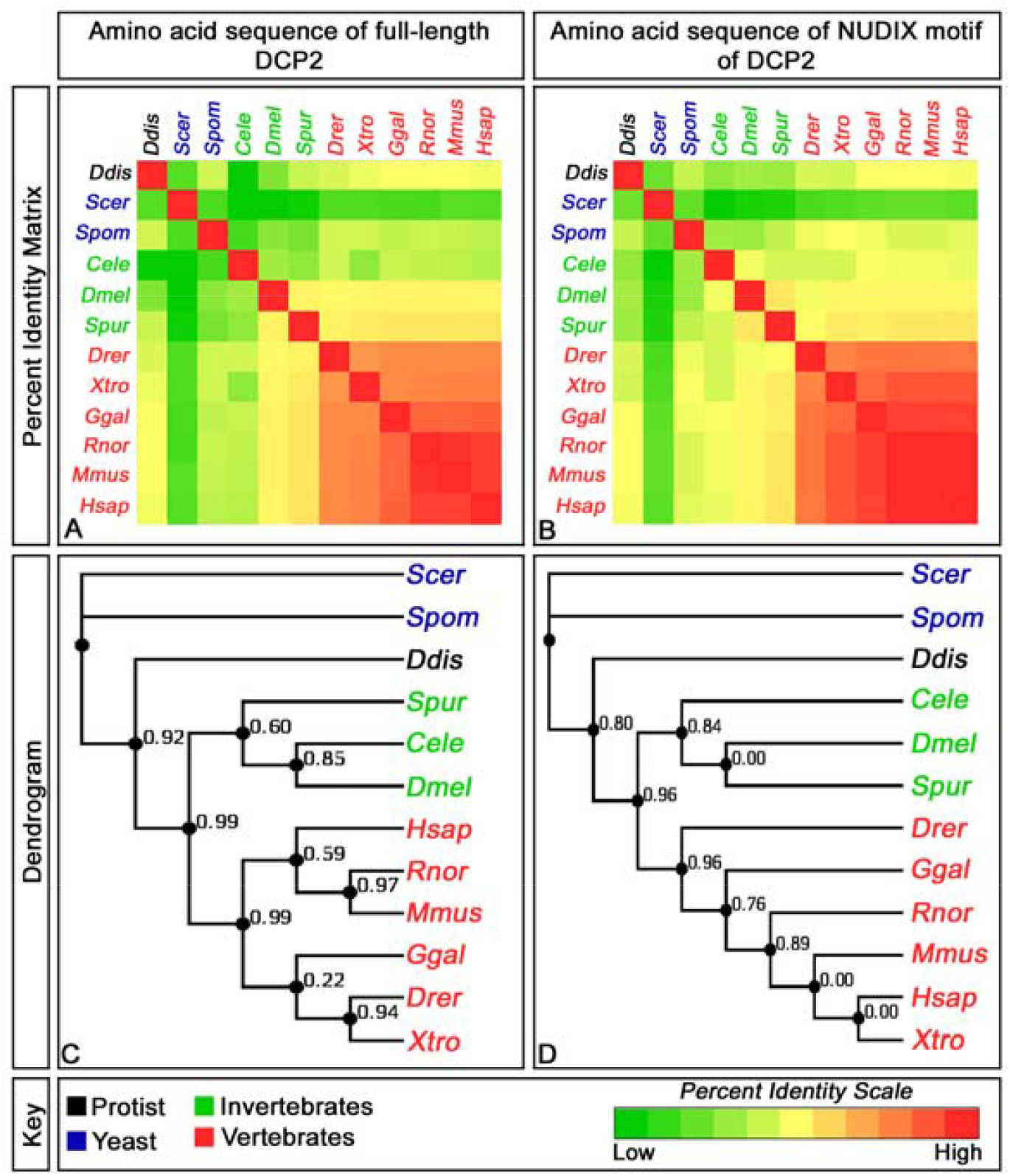
Sequence identity and evolutionary relationships of DCP2 orthologs across the phylogenetic tree. A shows the sequence similarity of the complete linear sequence of DCP2 across the different phyla while B shows the same for the sequence of the NUDIX motif in these orthologs. C and D show the respective dendrograms derived from the above percent identities in A and B respectively.

On inspection of the molecular phylogenetic relationships among the orthologs through Maximum Likelihood method, the yeasts harbor the oldest orthologs, while the *Xenopus* ortholog is the youngest. The vertebrate orthologs appear to have originated from a common ancestral form which the *C. elegans* (annelid), *D. melanogaster* (insect) and *S. purpuratus* (echinoderm) orthologs evolved.

Thus, the vertebrate orthologs show a higher degree of molecular conservation than the invertebrate members.

#### b. Among *Drosophila* sp., the *melanogaster* DCP2 ortholog is closest to that harbored by the *repleta* sub-group

Following the analysis across the different phyla of the evolutionary tree, the analysis was extended to the twelve species of *Drosophila*, whose genomic data was available on FlyBase. These species belonged to six different subgroups *viz*., *melanogaster* (*D. melanogaster, D. ananassae, D. erecta, D. yakuba, D. simulans* and *D. sechellia*), *obscura* (*D. persimilis* and *D. pseudoobscura pseudoobscura*), *virilis* (*D. virilis*), *repleta* (*D. mojavensis*), *Hawaiian* (*D. grimshawi*) and *willistoni* (*D. willistoni*) and were analysed similarly for the amino acid sequence composition of DCP2. *D. grimshawi* (*Hawaiian* species) was observed to harbor the longest ortholog (926 residues) while that harbored by *D. persimilis* (*obscura* subgroup) is the shortest (570 residues). The length of the N-terminus to the Box A domain is fairly uniform across the species except for *D. persimilis*, which has only one (1) amino acid residue preceding its Box A domain. Also, the Box A domain is composed of only 77 residues and starts with Glutamic acid (E), whereas in all the other species, it is composed of ∼93-95 residues and starts with Aspartic acid (D). *D. willistoni* however employs 115 residues to generate the same domain, but the pioneering (Aspartic acid; D) and the terminal (Tyrosine; Y) residues are the same as others. The tripeptide spacer sequence, *viz*., Lysine (K) –Leucine (L) – Serine (S), is conserved in all the species except that the central residue in the two species of the *obscura* subgroup is Methionine (M). The NUDIX domain is highly conserved with constant length (150 residues) and starts with Valine (V) and terminates in Isoleucine (I) (**Table 2 and Figure 3**).

**Table 2:**
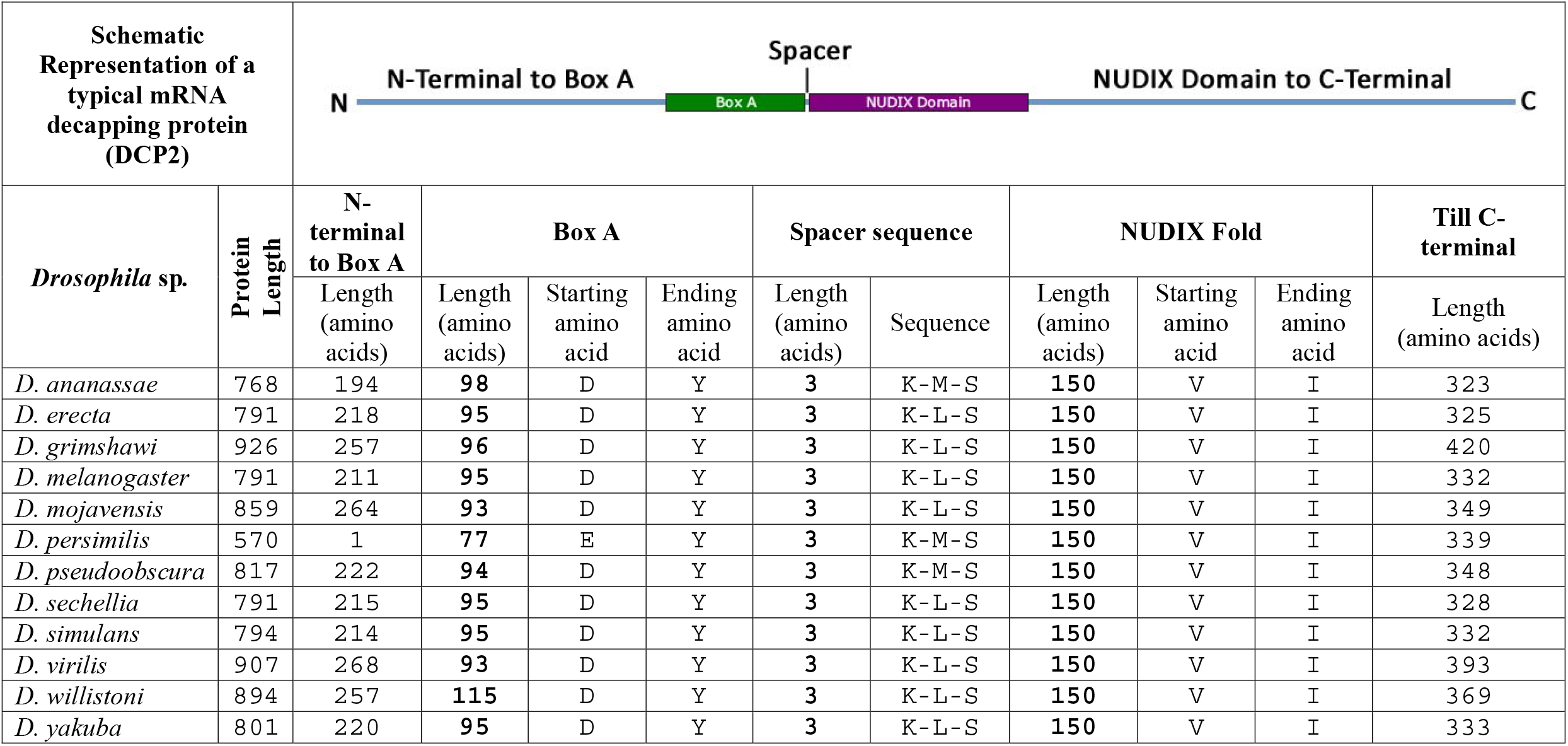
Table showing the DCP2 orthologs in sibling species of *Drosophila melanogaster*. The NUDIX domain is highly conserved with constant length and sequence whereas the Box A domain varies in size but almost always starts with Aspartic acid (D), except for *Drosophila persimilis* which starts with Glutamic acid (E), and ends with Tyrosine (Y). The Box A domain is followed by the spacer sequence towards the C-terminal, which is three amino acids long and begins with Lysine (K) and ends with Serine (S). Notably, *D. persimilis* has the shortest protein, with only 570 residues and has the shortest Box A domain as well while *D. grimshawi* has the longest protein with 926 residues, but the signature domains conform to the sizes observed in others. *D. willistoni* on the other hand, has the most number of residues composing the Box A domain.

| Schematic Representation of a typical mRNA decapping protein (DCP2) |  |  |  |  |  |  |  |  |  |  |  |
| --- | --- | --- | --- | --- | --- | --- | --- | --- | --- | --- | --- |
| *Drosophila* sp. | Protein Length | N-terminal to Box A | Box A |  |  | Spacer sequence |  | NUDIX Fold |  |  | Till C-terminal |
|  |  | Length (amino acids) | Length (amino acids) | Starting amino acid | Ending amino acid | Length (amino acids) | Sequence | Length (amino acids) | Starting amino acid | Ending amino acid | Length (amino acids) |
| *D. ananassae* | 768 | 194 | 98 | D | Y | 3 | K-M-S | 150 | V | I | 323 |
| *D. erecta* | 791 | 218 | 95 | D | Y | 3 | K-L-S | 150 | V | I | 325 |
| *D. grimshawi* | 926 | 257 | 96 | D | Y | 3 | K-L-S | 150 | V | I | 420 |
| *D. melanogaster* | 791 | 211 | 95 | D | Y | 3 | K-L-S | 150 | V | I | 332 |
| *D. mojavensis* | 859 | 264 | 93 | D | Y | 3 | K-L-S | 150 | V | I | 349 |
| *D. persimilis* | 570 | 1 | 77 | E | Y | 3 | K-M-S | 150 | V | I | 339 |
| *D. pseudoobscura* | 817 | 222 | 94 | D | Y | 3 | K-M-S | 150 | V | I | 348 |
| *D. sechellia* | 791 | 215 | 95 | D | Y | 3 | K-L-S | 150 | V | I | 328 |
| *D. simulans* | 794 | 214 | 95 | D | Y | 3 | K-L-S | 150 | V | I | 332 |
| *D. virilis* | 907 | 268 | 93 | D | Y | 3 | K-L-S | 150 | V | I | 393 |
| *D. willistoni* | 894 | 257 | 115 | D | Y | 3 | K-L-S | 150 | V | I | 369 |
| *D. yakuba* | 801 | 220 | 95 | D | Y | 3 | K-L-S | 150 | V | I | 333 |

**Figure 3:**
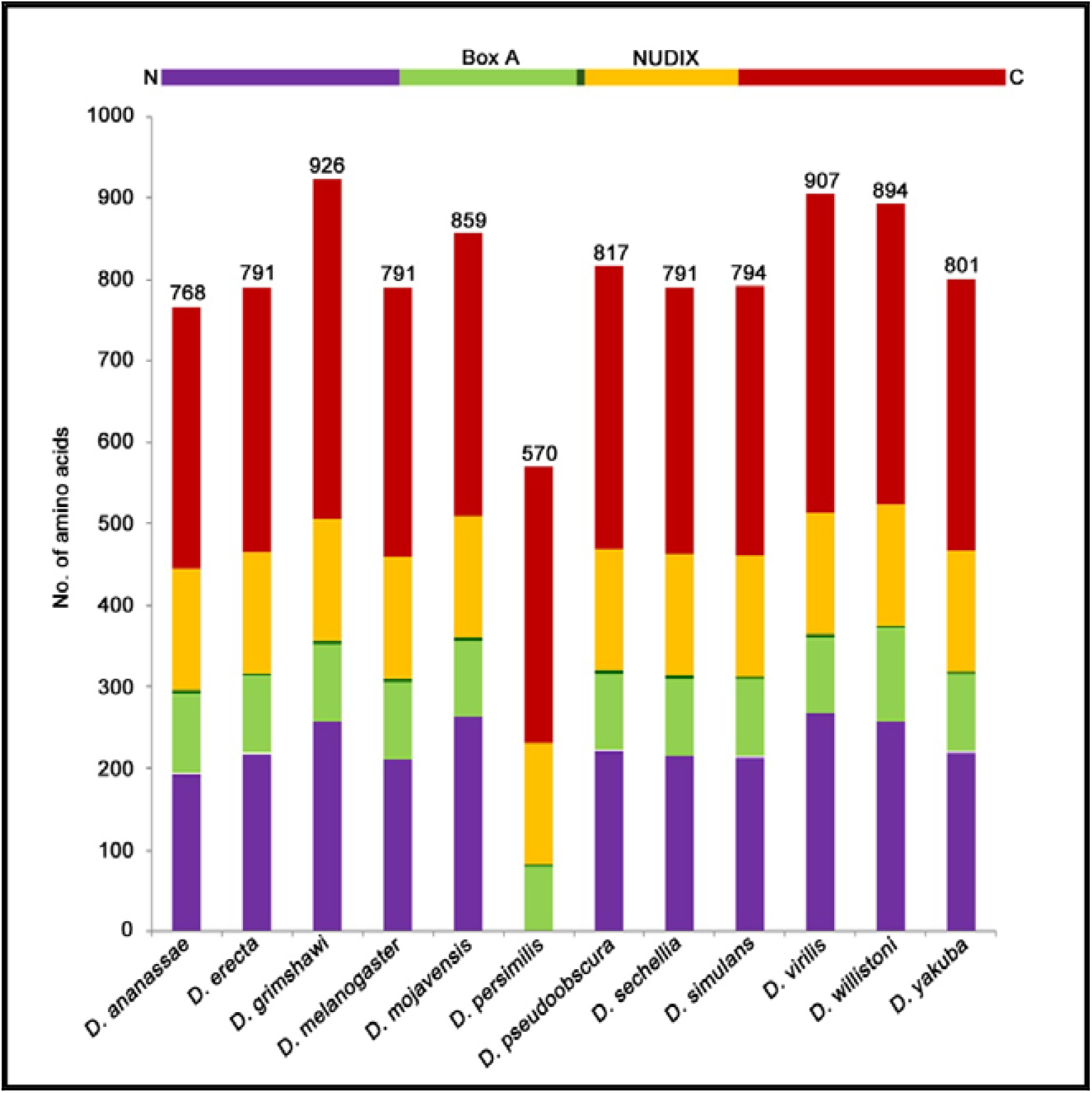
Graphical representation of the different regions of the DCP2 orthologs in sibling species of *Drosophila melanogaster*. *D. grimshawi* (*Hawaiian* species) harbours the longest ortholog (926 residues) while that harbored by *D. persimilis* (*obscura* subgroup) is the shortest (570 residues). The length of the N-terminus to the Box A domain is fairly uniform across the species except for *D. persimilis*, which has only one (1) amino acid residue preceding its Box A domain). The NUDIX domain is highly conserved with constant length (150 residues).

Closer analyses of the complete DCP2 sequence and the sequence of the NUDIX motif harbored, through percentage identity matrices (**Figure 4 A and B**), shows that the *melanogaster* orthologs are more conserved in either case as compared to their siblings. Strikingly, although *D. ananassae* belongs to the same subgroup, it shows the least similarity with the other species of the subgroup. Moreover, the *D. persimilis* ortholog shows better homology with the species of the *melanogaster* subgroup analysed. However, in all the species considered here, the sequence of the NUDIX domain shows a better index of homology as compared to the sequence of the entire DCP2 protein, and the observed differences in the total protein sequence is majorly attributed to the differences in the N- and C-terminal sequences. The only exceptions are observed in the *virilis* (*D. virilis*), *repleta* (*D. mojavensis*), *Hawaiian* (*D. grimshawi*) species, where the sequence of the complete protein ortholog is more similar than that of the NUDIX domain.

**Figure 4:**
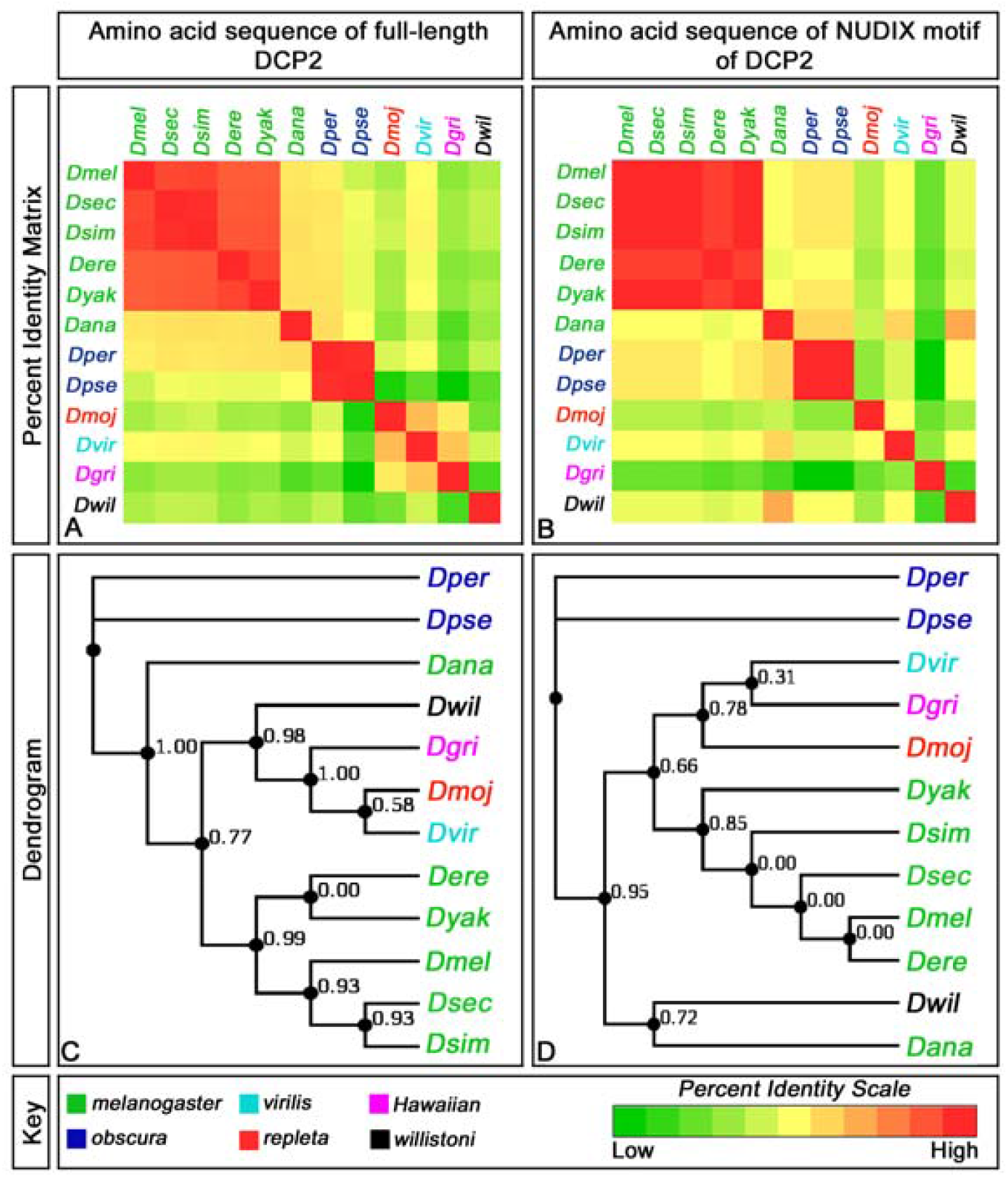
Sequence identity and evolutionary relationships of DCP2 orthologs in sibling species of *Drosophila melanogaster*. A shows the sequence similarity of the complete linear sequence of DCP2 across the different species of *Drosophila*, belonging to six different subgroups while B shows the same for the sequence of the NUDIX motif in these orthologs. C and D show the respective dendrograms derived from the above percent identities in A and B respectively.

On inspection of the molecular phylogenetic relationships among the orthologs through Maximum Likelihood method, the *obscura* subgroup harbors the oldest orthologs. The *melanogaster* orthologs appear to have originated from a common ancestral form which the *virilis* (*D. virilis*), *repleta* (*D. mojavensis*), *Hawaiian* (*D. grimshawi*) orthologs evolved. Plausibly, the *melanogaster* ortholog is closer to that harbored by the *repleta* subgroup.

Hence, from the analyses of the different representative species across the phylogenetic tree, the sequence of the NUDIX domain is evidently more conserved than that of the rest of the protein.

### *In silico* predictions of the DmDCP2 structure identify distinct topological paradigms in the secondary and tertiary structures

Structural features of proteins provide newer insights into their functional potential by throwing light on the mechano-dynamic properties, plausible interactome and the possible functional diversity. Determination of structure of a protein involves multiple stages and is accomplished through standard biophysical approaches, the most popular ones involving as analyses of x-ray diffraction (XRD) patterns by crystals of the protein or nuclear magnetic resonance (NMR) spectroscopic analyses of the protein in solution (Alberts, 2002). However, both of these methods require the protein to be over-expressed, isolated and purified through rigorous molecular biological protocols in the wet lab. With the advent of *in silico* modeling platforms, which require only the primary sequence of the protein, it is possible to generate putative models to identify and analyse the putative structural features of any given protein. In the present analysis, instead of using homology based modeling approach, the sequence was first assessed for its secondary structure and then a threading based approach was employed to generate the tertiary structure using the I-TASSER platform (Zhang, 2008; Roy et al., 2010), which is an online server and generates protein structures by iterative fragment assembly simulations.

#### a. Secondary structure of DmDCP2

Prediction of secondary structure of the *Drosophila* mRNA decapping protein 2 using PSIPRED (**Figure 5**) shows that the sequence is conducive for the formation of a number of helices, connected by random coils and that very few regions engage in the formation of beta strands. Two sets of sequences, depicting the two classic motifs of DCP2, are underlined. Amino acids 212-306, underlined green and consist solely of alpha helices, form the Box A domain (pfam05026), while the amino acids 310-459, underlined purple and consist of alpha helices interspersed with beta strands, form the classic NUDIX (<u>**Nu**</u>cleoside <u>**di**</u>phosphate linked to moiety <u>**X**</u>; cd03672) domain which catalyses the removal of the methylguanosine cap from the five prime end of mRNAs. Most notably, the only beta strands in the entire protein are the ones forming the NUDIX Motif.

**Figure 5:**
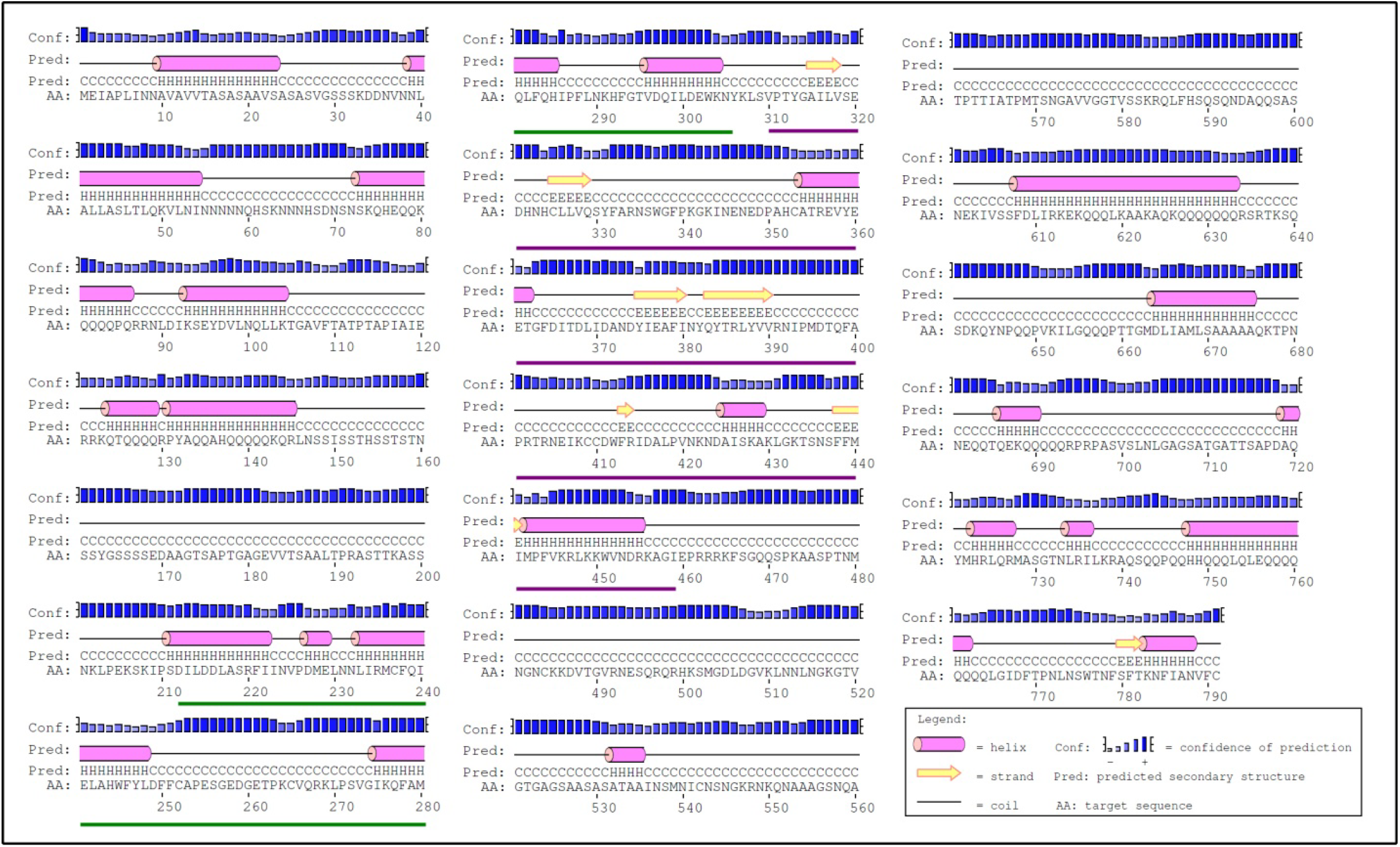
Predicted secondary structure of DmDCP2-PA. Predicted secondary structure of the *Drosophila* mRNA decapping protein 2 shows that the sequence is conducive for the formation of a number of helices connected by random coils. Two sets of sequences, depicting the two classic motifs of DCP2, are underlined. Amino acids 212-306, underlined **green** and consist solely of alpha helices, form the Box A domain while the amino acids 310-459, underlined **purple** and consist of alpha helices interspersed with beta strands constitute the NUDIX domain. The only beta strands in the entire protein are the ones forming the NUDIX Motif.

#### b. The tertiary structure of DmDCP2 shows an evolutionarily conserved topology of the NUDIX domain

The tertiary structure of DmDCP2 (DCP2-PA; 791 residues) shows that most of the protein engages in the formation of random coils which are interspersed with short helices and the only beta strands are found in the NUDIX domain (**Figure 6A**). While the Box A domain is an all helical structure, the NUDIX domain is a compact structure in the tree-dimensional space, being composed of four (4) alpha helices (*viz*., α_1N-4N_) and six (6) beta strands (*viz*., _β1N-4N_) (**Figure 6 A; inset**). The N-terminal regulatory domain (RD; Wurm and Sprangers, 2019) is mostly comprised of tightly packed random coils whereas the C-terminal intrinsically disordered region (IDR; Wurm and Sprangers, 2019) consists of random coils and small helices packed loosely, similar to the human ortholog. To assess the backbone confirmation, the Phi (φ) /Psi (ψ) angles were analysed using the Ramachandran plot (Ramachandran *et al*., 1963), which showed 55.6% (439/791) residues to reside in the *favourable region* and 23.2% (183/791) residues to reside in the *allowed region*, while only 21.2% (167/791) residues were found to be in the *outlier region* (**Figure 7**), which showed that the model may be quite close to that obtained by wet lab approaches. The protein binds to guanosine triphosphate (GTP), which is coordinated by the Asp_396_, Gln_398_, Ala_400_ and the Arg_402_ residues (**Figure 8 A**), all of which belong to the NUDIX domain. A nitrogen atom (N2) from the GTP moiety also forms a hydrogen bond with the Ala_400_ residue. DCP2 requires magnesium ions (Mg^2+^) ions for its activity and in the model obtained, the Mg^2+^ ion is found to be coordinated by Glu_357_ and Arg_404_ residues (**Figure 8 B**). On comparing the structure with the crystal structures of the yeast (PDB ID: 5J3Y; Chain A) and human (PDB ID: 5QPC; Chain A), orthologs, the topology of the NUDIX domain of the generated model was found to be in complete alignment with that of the crystal structures (**Figure 8 B and C**), thereby revealing an evolutionarily conserved topology of the NUDIX domain despite sequence diversity.

**Figure 6:**
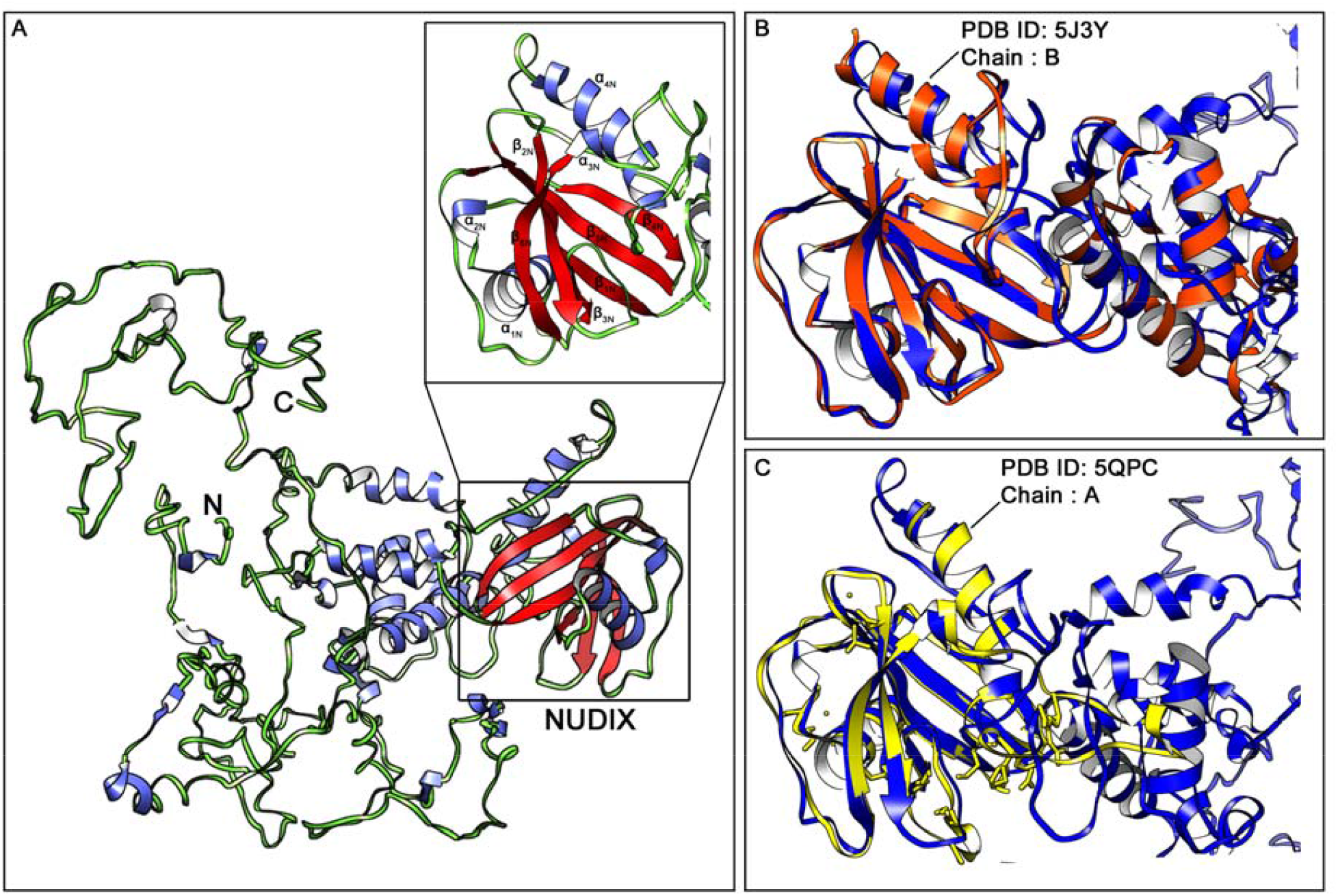
Predicted tertiary structure of DmDCP2-PA. Predicted tertiary structure of the *Drosophila* mRNA decapping protein 2 shows that most of the protein engages in the formation of **random coils** interspersed with short **helices**. The only **beta strands** are found in the NUDIX domain (A). While the Box A domain is an all helical structure, the NUDIX domain (A; inset) is a compact structure in the three-dimensional space, being composed of four (4) alpha helices (*viz*., α_1N-4N_) and six (6) beta strands (*viz*., _β1N-4N_). The N-terminal RD is mostly comprised of tightly packed random coils whereas the C-terminal IDR consists of random coils and small helices packed loosely, similar to the human ortholog. Structural comparison *vis-à-vis* alignment with the crystal structures of the yeast (B) and human (C) orthologs show the topology of the NUDIX domain of the generated model to be in complete alignment with that of the crystal structures.

**Figure 7:**
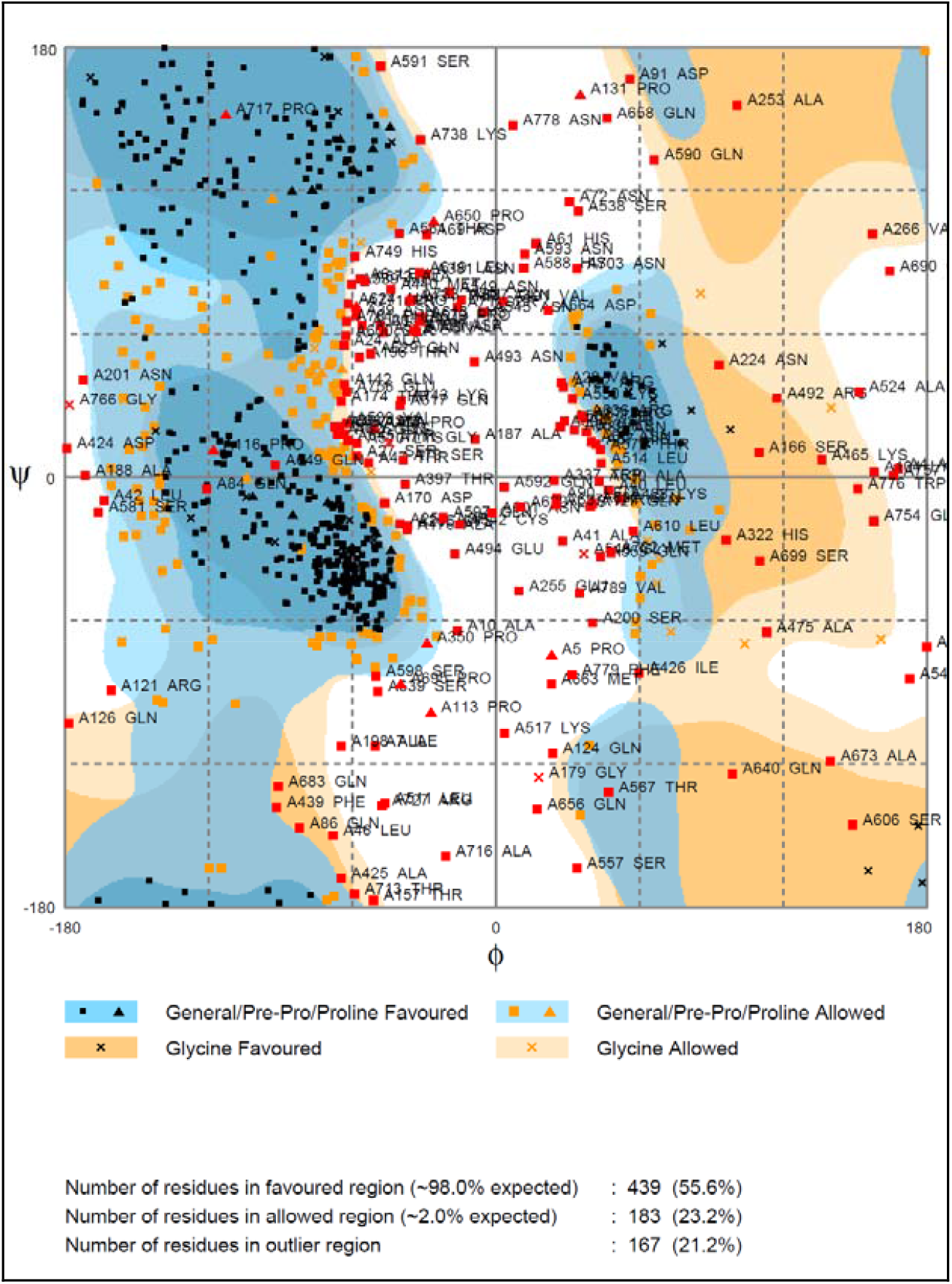
Ramachandran plot of the predicted model of DmDCP2. Analysis of the backbone conformation, *i*.*e*., the Phi (φ) /Psi (ψ) angles using the Ramachandran plot showed 55.6% (439/791) residues to reside in the *favourable region* and 23.2% (183/791) residues to reside in the *allowed region*, while only 21.2% (167/791) residues were found to be in the *outlier region*, implying the model to be quite close to that obtained by wet lab approaches.

**Figure 8:**
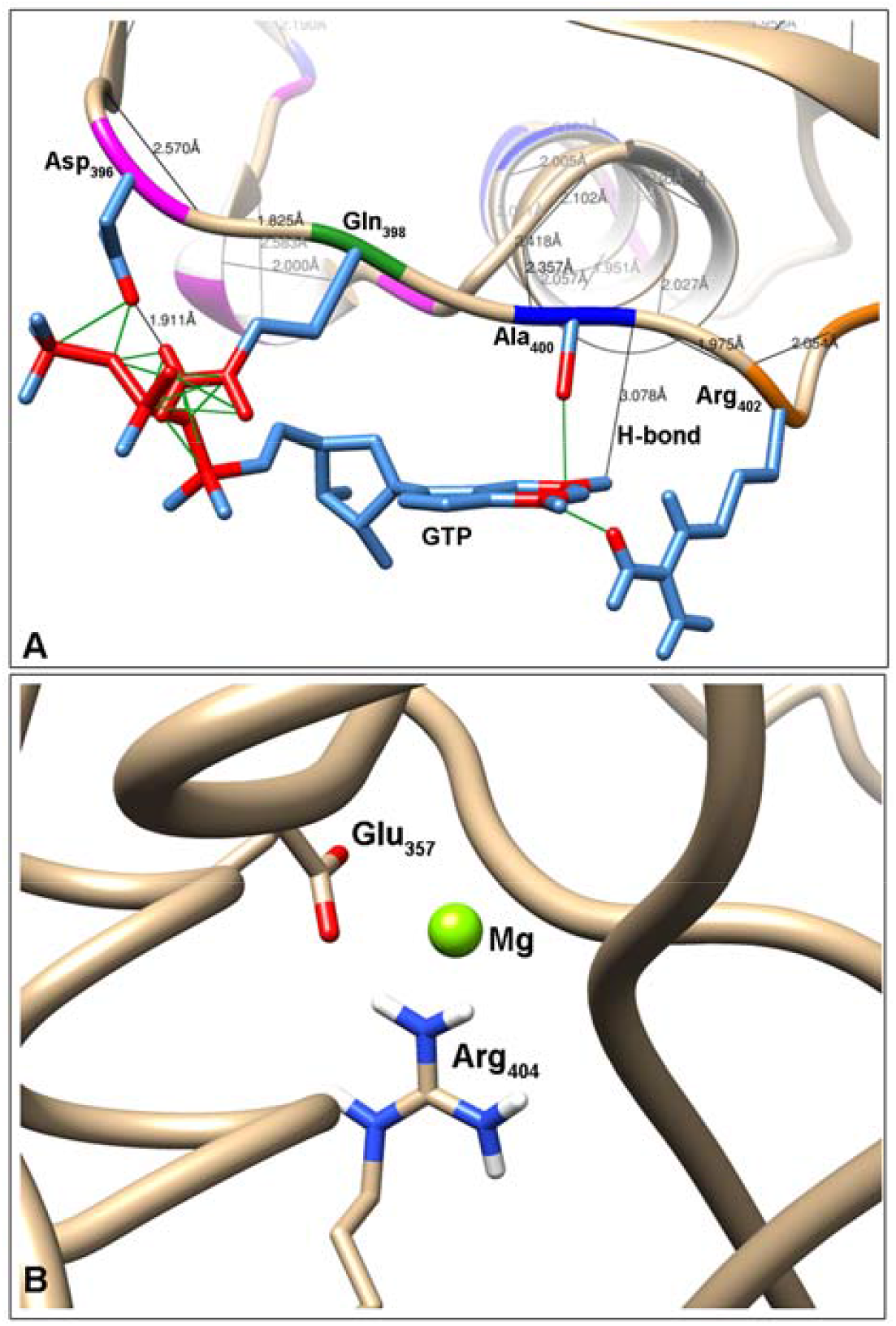
Coordination of DmDCP2 to GTP and Mg^2+^. The GTP is coordinated by the Asp_396_, Gln_398_, Ala_400_ and the Arg_402_ residues (A), all of which belong to the NUDIX domain. A nitrogen atom (N2) from the GTP moiety also forms a hydrogen bond with the Ala_400_ residue. DCP2 requires magnesium ions (Mg^2+^) ions for its activity and in the model obtained, the Mg^2+^ ion is found to be coordinated by Glu_357_ and Arg_404_ residues (B).

### NUDIX proteins *vis-à-vis* DCP2 paralogs in *D. melanogaster* show topological conservation of the NUDIX domain despite sequence divergence

Although the NUDIX proteins consist primarily of pyrophosphohydrolases (Bessman et al, 1996), they are involved in multiple functions in the cell and some of the NUDIX proteins are non-enzymatic in nature and function as modulators of cellular function by interacting with other proteins or are modulators of transcription (Srouji et al, 2017). Despite availability of information pertaining to the kinetics, evolution or dynamics of NUDIX proteins through *in silico* or wet molecular approaches, the identification and understanding of other NUDIX proteins in *Drosophila* is still in its infancy and remains unexplored. Hence, in order to identify the other NUDIX proteins in *D. melanogaster*, the amino acid sequence of the DCP2 NUDIX domain was used as “bait” in a homology search to fish out other NUDIX proteins. Since, all these proteins bear the same functional catalytic domain – the NUDIX domain, these may be referred to, as the paralogs of DCP2.

These proteins were assessed for the conservation of sequence of the complete protein and then for the NUDIX domain only, following which, the molecular phylogenetic relationships were also identified. The proteins were then modelled to deduce their tertiary structure, which were then aligned individually with the generated structure of DmDCP2 to identify regions of structural homology.

#### a. NUDIX proteins in *D. melanogaster* show extremely low sequence conservation

Through a homology search in the FlyBase, twelve (12) NUDIX proteins were identified besides DCP2, out of which eight (8; including DCP2) had their NUDIX sequence annotated. **Table 3** shows the list of the NUDIX proteins identified in the homology search, their sizes, the length of the NUDIX domain including the pioneering and terminal residues of it, the presence of other domains in the rest of the protein chain and their subcellular location. All the proteins identified are hydrolases with different substrate specificity and thus perform different functions in the cell. Out of the identified proteins, one protein, CG2091 is the putative Scavenger decapping enzyme (DcpS). Most notably, except DCP2, all the other proteins are shorter in length and smaller in size.

**Table 3:** Table showing the NUDIX proteins in *Drosophila melanogaster*. Besides DCP2, twelve (12) NUDIX proteins were identified in the homology search, out of which eight (8; including DCP2) had their NUDIX sequence annotated. Some of their features are enlisted below.

| CG Identifier (FlyBase) | Name | Function | Size (amino acids) | NUDIX |  |  | Other domain Length (amino acids) | Cellular location |
| --- | --- | --- | --- | --- | --- | --- | --- | --- |
|  |  |  |  | Length (amino acids) | Starting amino acid | Ending amino acid |  |  |
| CG6169 | DCP2 | m<sup>7</sup>G(5')pppN diphosphatase; Mn<sup>2+</sup> binding; RNA binding | 791 | 150 | Val (310) | Ile (459) | Box A (95) (202-306) | Cytoplasm |
| CG2091 | DcpS | m<sup>7</sup>G(5')pppN diphosphatase; RNA 7-methylguanosine cap binding | 374 | 109 | Asp (16) | Ser (124) | - | Nucleus; Cytoplasm |
| CG31713 | Apf | Bis(5'-nucleosyl)-tetraphosphatase (asymmetrical) (3.6.1.17) | 142 | 113 | Ala (4) | Lys (116) | - | - |
| CG6391 | Aps | Endopolyphosphatase (3.6.1.10; Diphosphoinositol-polyphosphate diphosphatase (3.6.1.52) | 177 | 119 | Arg (18) | Gln (136) | - | - |
| CG8128 | - | ADP-ribose diphosphatase (3.6.1.13); Nucleotide diphosphatase (3.6.1.9); NAD binding | 330 | 122 | Gly (163) | Val (284) | Pre-NUDIX (80) (67-146) | - |
| CG12567 | - | 8-oxo-dGDP phosphatase | 349 | 105 | Ile (154) | Cys (258) | DUF4743 (123) (14-136) | - |
| CG42813 | - | UDP-sugar diphosphatase (3.6.1.45) | 212 | 145 | Val (40) | Gln (184) | - | - |
| CG10898 | - | Hydrolase | 340 | 101 | Val (61) | Arg (161) | - | - |
| CG10194 | - | Hydrolase | 351 | NUDIX sequence not annotated |  |  | - | - |
| CG10195 | - | Hydrolase | 361 |  |  |  | - | - |
| CG11095 | - | Hydrolase | 283 |  |  |  | - | - |
| CG18094 | - | Hydrolase | 360 |  |  |  | - | - |
| CG42814 | - | UDP-sugar diphosphatase (3.6.1.45) | 237 |  |  |  | - | - |

Closer analyses of the complete sequence of the protein and the sequence of the NUDIX motif harbored, through percentage identity matrices (**Figure 9 A and B**), shows that the proteins show extremely low sequence similarity among themselves. Strikingly, DCP2 shows very low homology with the scavenger decapping enzyme, DcpS, despite functional similarity. In the PIM, CG8128, which is a nucleotide diphosphatase *vis-à-vis* ADP-ribose diphosphatase, and CG10898, which is a hydrolase with uncharacterized function are the only pair of proteins which show a better index of sequence similarity as compared to paired analysis of other proteins. Although all the proteins bear the NUDIX motif, the sequence of the motif varies among them and shows extremely low degree of conservation.

**Figure 9:**
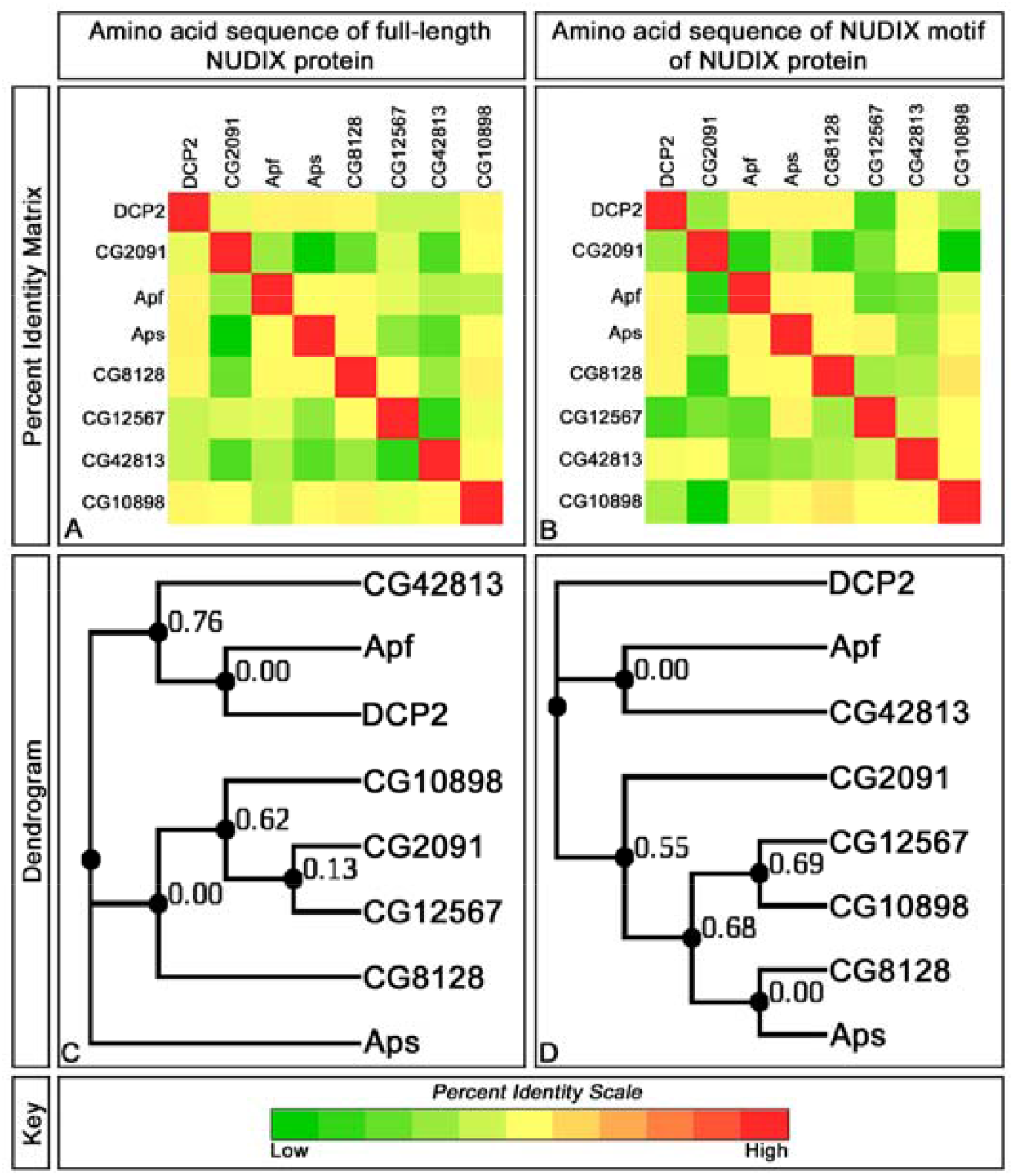
Sequence identity and evolutionary relationships of DCP2 paralogs in *Drosophila melanogaster*. A shows the sequence similarity of the complete linear sequence of the NUDIX proteins while B shows the same for the sequence of the NUDIX motif in these orthologs. C and D show the respective dendrograms. Strikingly, DCP2 shows very low similarity DCPS, despite functional similarity. Most notably, CG8128 and CG10898 have a better index of sequence similarity as compared to other proteins. DCP2 shares a closer phylogenetic relationship with Apf as compared to other proteins and seems to have evolved from a common ancestral protein with Apf.

On inspection of the molecular phylogenetic relationships among the orthologs through Maximum Likelihood method, it seems that the NUDIX motifs and the sequence of the complete protein *per se* of DCP2 (a diphosphatase), Apf (a tetraphosphatase) and CG42813 (a diphosphatase) have plausibly evolved from a common ancestral sequence, which diverged across time to give way to the functional divergence observed. Also, DcpS (decapping enzyme; diphosphatase) presumably shares a closer phylogenetic relationship with CG12567 (8-oxo-dGTP-phosphatase) instead of DCP2 and is a more recently evolved protein unlike DCP2.

#### b. The topology of the NUDIX domain is conserved despite lack of sequence conservation

On modeling the NUDIX proteins *in silico* (**Figure 10**), all the proteins showed their NUDIX domains being composed of alpha helices and beta strands, similar to that observed in DCP2, but all the proteins had beta strands in other regions of the protein sequence as well, unlike DCP2. Moreover, they lack the extensive stretches of random coils and occupy a smaller volume in space. However, on aligning them to the structure of DCP2 so generated, the tertiary structure of the NUDIX motif of all the proteins, except DcpS and CG8128 were similar to each other and to that of DCP2 (**Figure 11**), reflecting an evolutionary conservation of the structure of the NUDIX domain despite lack of sequence similarity *vis-à-vis* conservation. Although DCP2 and DcpS share a functional similarity, they do not show similarities in either sequence or structure, even in the functional domain which confers the functional identity and similarity to them.

**Figure 10:**
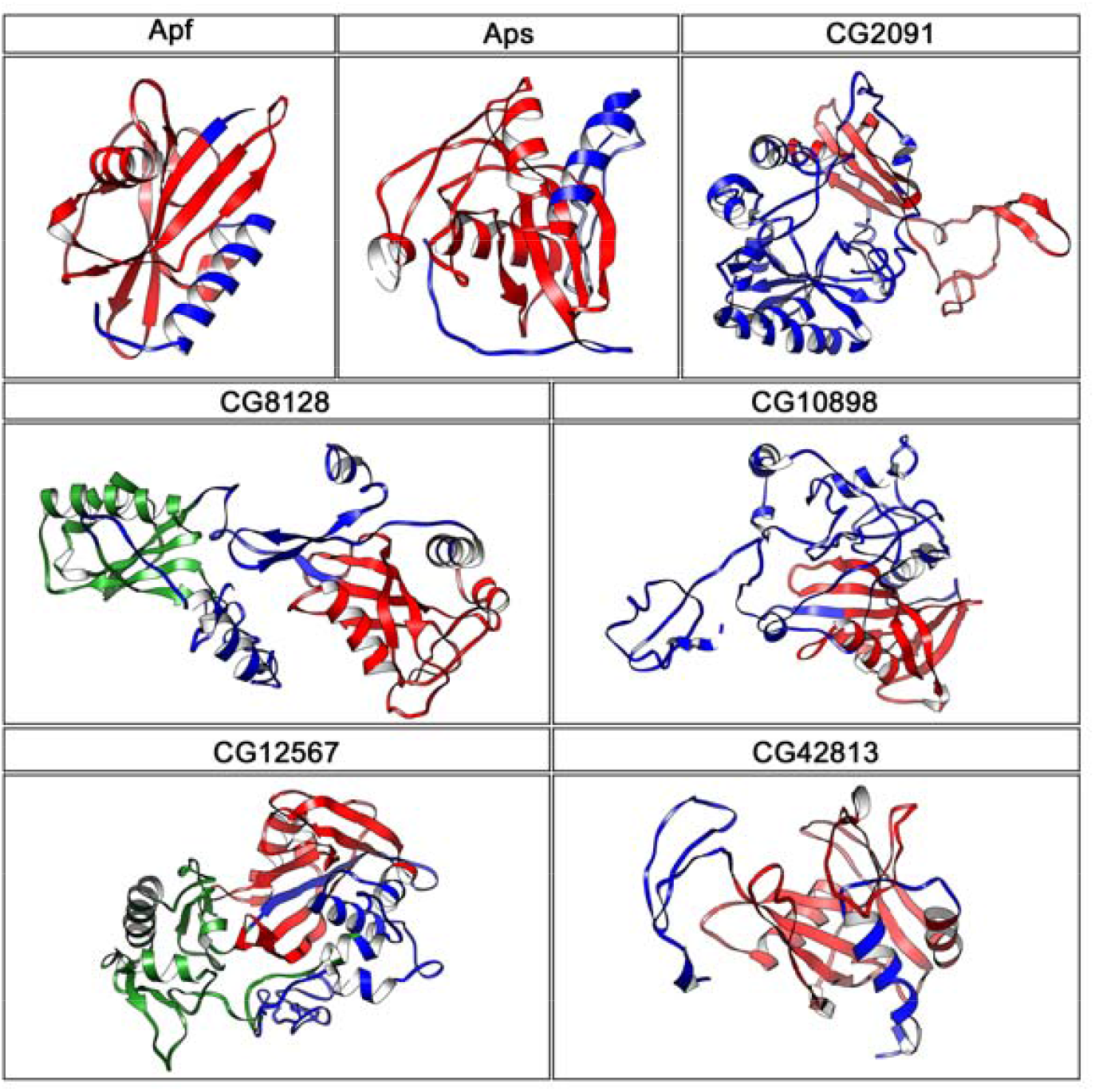
Predicted tertiary structures of the NUDIX proteins in *D. melanogaster*. All the proteins have NUDIX motif composed of alpha helices and beta sheets. Most strikingly, the other NUDIX proteins are composed of fewer amino acids as compared to DCP2 and have beta sheets in non-NUDIX regions as well unlike DCP2 which harbours beta sheets only in the NUDIX domain. Also, they lack the extensive stretches of random coils and occupy smaller volume in space. The **NUDIX** domain is painted in **red**, while the **alpha helices** and **beta strands** are depicted in **green** and **blue** respectively.

**Figure 11:**
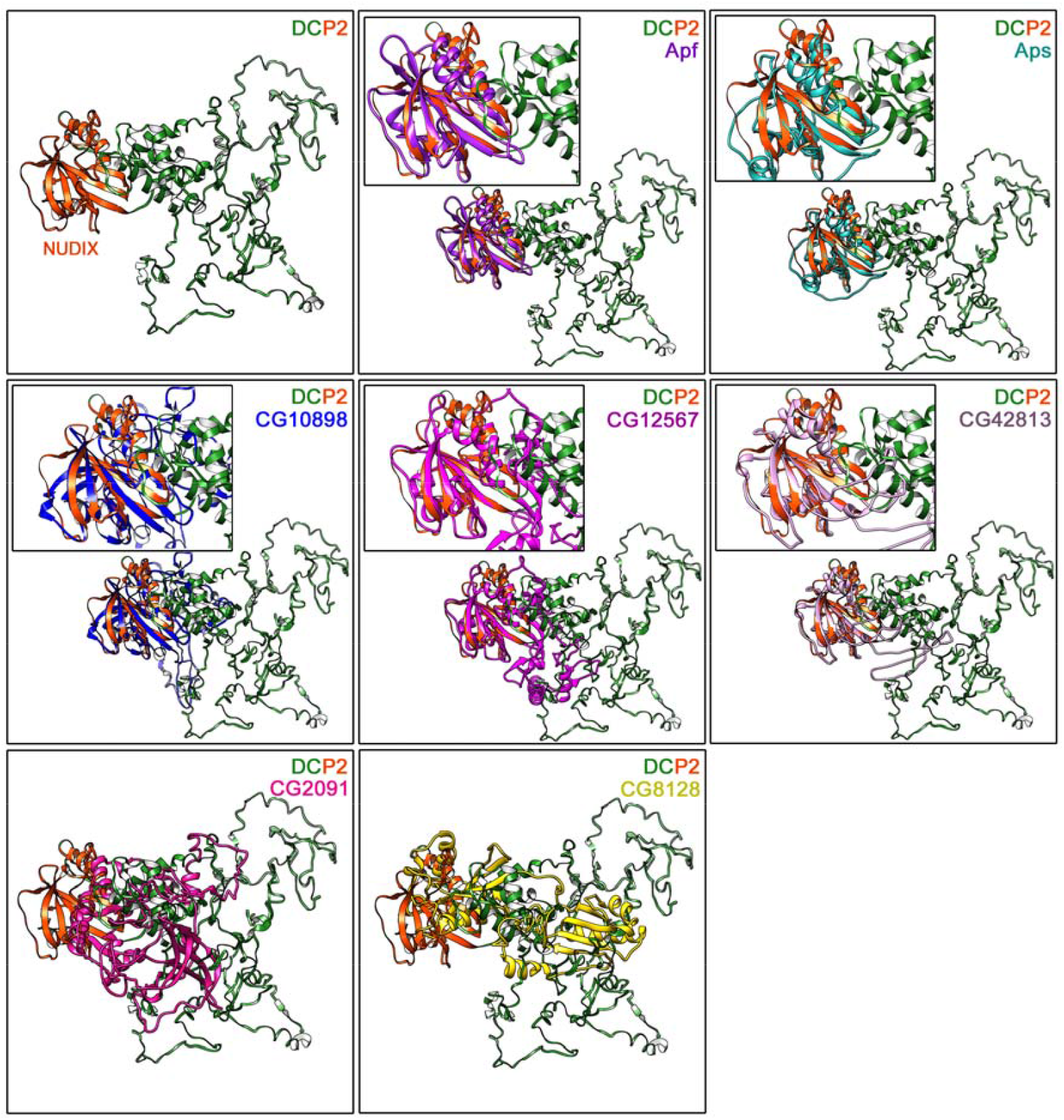
The topology of the NUDIX motif is conserved beyond sequence dissimilarity. The structure of DCP2 is represented in the first model, wherein the **NUDIX** motif is painted in **orange**, and the remaining chain in **green**. For each of the other models, the structural overlap of the NUDIX motif of the protein with that of the DCP2 NUDIX motif is shown in the inset. Notably, the tertiary structure of the NUDIX motif of almost all the proteins is similar to that of DCP2, despite dissimilarity in the linear sequence. NUDIX motifs of CG2091 (DCPS) and CG8128 do not overlap with each other or with that of any other NUDIX protein.

## Conclusion

The present endeavor is an effort to identify and discern the extent of conservation of the amino acid sequence of the mRNA decapping protein 2, DCP2, which is NUDIX protein, in different species across the phylogenetic tree and in the sibling species of *Drosophila melanogaster*, which occupy the same taxon in the tree, and to identify the similar relationships of DCP2 with its paralogs in *D. melanogaster*. The study shows that while the sequence of the NUDIX domain is much more conserved than the rest of the protein sequence across species, the complete amino acid sequence of DCP2 shows increasing degree of conservation as we ascend the evolutionary tree, with the vertebrate orthologs emerging from a common ancestral isoform. Interestingly, although functionally conserved, a gradual decrease in the size of the protein in visible parallel to the ascent in the tree, and the decrease occurs primarily in the sequences beyond the NUDIX motif, *i*.*e*., at the N- and C-termini. On modeling the structure of DmDCP2, the tertiary structure of the NUDIX motif reflects an evolutionary conservation of topology, showing structural homology *vis-à-vis* conservation with the lowest (yeast) and the highest (human) DCP2 structures despite sequence divergence along evolution, which is in agreement with the fact that the domain architecture of orthologous proteins is always conserved (Forslund et al, 2011). An inspection of the other NUDIX proteins in *D. melanogaster* with reference to the conservation of their sequence and structure reinforces the observation that despite the evolution of protein sequence by mutations, both locally and/or globally, which may have given rise to their functional divergence, the structure of functional domains is far more conserved than sequence (Illergard et al, 2009).

Hence, the present observations provide quantitative and structural bases for the observed evolutionary conservation of DCP2 across the diverse phyla and also, identify and reinforce the structural conservation of the NUDIX family in *D. melanogaster*.

## Author Contributions

RK, conceptualization, resources, methodology, investigation, data curation, formal analysis and interpretation, writing the manuscript. JKR, supervision.

## Conflict of Interest

The authors declare no conflict of interest.

## Notes

### Competing Interest Statement

The authors have declared no competing interest.

